# RRP7A links primary microcephaly to radial glial cells and dysfunction of ribosomal biogenesis, neurogenesis and ciliary resorption

**DOI:** 10.1101/793877

**Authors:** Muhammad Farooq, Louise Lindbæk, Nicolai Krogh, Canan Doganli, Cecilie Keller, Maren Mönnich, Srinivasan Sakthivel, Yuan Mang, Ambrin Fatima, Vivi Søgaard Andersen, Muhammad. S. Hussain, Hans Eiberg, Lars Hansen, Klaus Wilbrandt Kjaer, Jay Gopalakrishnan, Lotte Bang Pedersen, Kjeld Møllgård, Henrik Nielsen, Shahid. M. Baig, Niels Tommerup, Søren Tvorup Christensen, Lars Allan Larsen

**Affiliations:** Department of Cellular and Molecular Medicine, University of Copenhagen, Copenhagen, Denmark; Department of Bioinformatics and Biotechnology, Government College University, Faisalabad, Pakistan; Department of Biology, University of Copenhagen, Copenhagen, Denmark; Human Molecular Genetics Laboratory; Health Biotechnology Division, National Institute for Biotechnology and Genetic Engineering PIEAS, Faisalabad, Pakistan; Institute of Biochemistry I, University of Cologne, Cologne, Germany; Cologne Center for Genomics and Center for Molecular Medicine Cologne, University of Cologne, Cologne, Germany; Institute of Human Genetics, Heinrich-Heine-University, Düsseldorf, Germany

**Keywords:** RRP7A, brain development, radial glial cells, microcephaly, MCPH, neurogenesis, cell cycle control, ribosome biogenesis, nucleoli, primary cilia, centrosome

## Abstract

Primary microcephaly (MCPH) is characterized by reduced brain size and intellectual disability^1^. The exact pathophysiological mechanism underlying MCPH remains to be elucidated, but dysfunction of neuronal progenitors in the developing neocortex plays a major role^1^. Using homozygosity mapping and whole exome sequencing, we identified a homozygous missense mutation (p.W155C) in *Ribosomal RNA Processing 7 Homolog A, RRP7A*, which segregated with MCPH in a consanguineous family with 10 affected individuals. RRP7A is expressed in neural stem cells/radial glial cells of the developing human forebrain, and targeted mutation of *Rrp7a* leads to defects in both neurogenesis and proliferation in a mouse stem cell model. RRP7A localizes to centrosomes, cilia and nucleoli, and patient-derived fibroblasts display defects in processing of ribosomal RNA, resorption of primary cilia and cell cycle progression. Finally, analysis of zebrafish embryos with loss-of-function mutation in *rrp7a* confirmed that RRP7A depletion causes reduced brain size, impaired neurogenesis and cell proliferation as well as defective ribosomal RNA processing. These findings provide novel insight into human brain development and MCPH.

## Main

During development of the human neocortex, neuroepithelial cells (NECs) multiply by symmetric cell division to create an initial pool of neural progenitors (NPCs), which differentiate into apical radial glial cells (RGCs) in the ventricular (VZ) and inner subventricular (ISVZ) zones as well as to basal RGCs at the outer subventricular zone (OSVZ)^2^. RGCs undergo a complex pattern of symmetric and asymmetric cell divisions and differentiation, which expands the NPC pool and produces post-mitotic neurons^3, 4^. Tight regulation of cell division and differentiation in NECs, RGCs and their progeny is important for proper NPC expansion and ultimately the number of cortical neurons^5^. This number is reduced in autosomal recessive primary microcephaly (MCPH; MIM #251200); a rare neurodevelopmental disorder characterized by congenital reduction in occipitofrontal circumference due to hypoplasia of the cerebral cortex causing reduction in brain volume and a simplified gyral pattern. MCPH patients display varying degree of non-progressive cognitive dysfunction. Additionally, most MCPH patients typically show sloping forehead, but no other facial or physical abnormalities^6^. Genetic analyses of consanguineous families have identified at least 18 MCPH genes that regulate RGC function and NPC expansion, and encode proteins essential for centrosome and cilium biogenesis and various functions in transcriptional regulation, DNA damage responses and mitosis^1^.

We ascertained a five-generation family in which ten individuals born to consanguineous couples presented with moderate to severe MCPH, varying degree of intellectual disabilities and impaired cognitive function; four affected individuals exhibited severe speech impairment, but none had seizures or epileptic symptoms (Fig. 1a,b; Supplementary Table 1). Cerebral MRI scans of one patient (V-14) showed decreased craniofacial ratio, slanting forehead and normal dimensions of the ventricular system (Fig. 1c). Volume loss was observed in the corpus callosum, especially in the anterior half. Linkage analysis identified two possible loss of heterozygosity (LOH) regions at chromosome 2q21.3 and 22q13.1-13.2. Further analysis using short tandem repeat and nucleotide polymorphism markers excluded linkage to 2q21.3 and confirmed linkage with a maximum LOD score Z=8.61 to a 2.5 Mb homozygous region at 22q13.1-13.2 (40,436,371-43,001,960 bp, hg38) (Fig. 1a, Supplementary Fig. 1a). This region contains 49 protein coding genes. Whole Exome Sequencing (WES) identified a very rare homozygous c.465G>C (p.W155C) missense mutation in *Ribosomal RNA Processing 7 Homolog A, RRP7A*, which is located within the linkage region (Fig. 1d, Supplementary Fig. 1b). Sanger sequencing confirmed that all affected family members were homozygous for the mutation, while parents of affected children were heterozygous carriers (Fig. 1a). The mutation was absent in 300 controls of Pakistani origin. The allele frequency of the mutation is 3.3E-05 in the South Asian population and the mutation is not present in 107,000 controls of other ethnicities (gnomad.broadinstitute.org). W155 in RRP7A is conserved from zebrafish to humans (Supplementary Fig. 1c), and *in silico* analysis of the effect of p.W155C mutation using the Combined Annotation Dependent Depletion (CADD) algorithm^7^ resulted in a scaled CADD score of 27.1, supporting that the mutation is pathogenic.

**Fig. 1:**
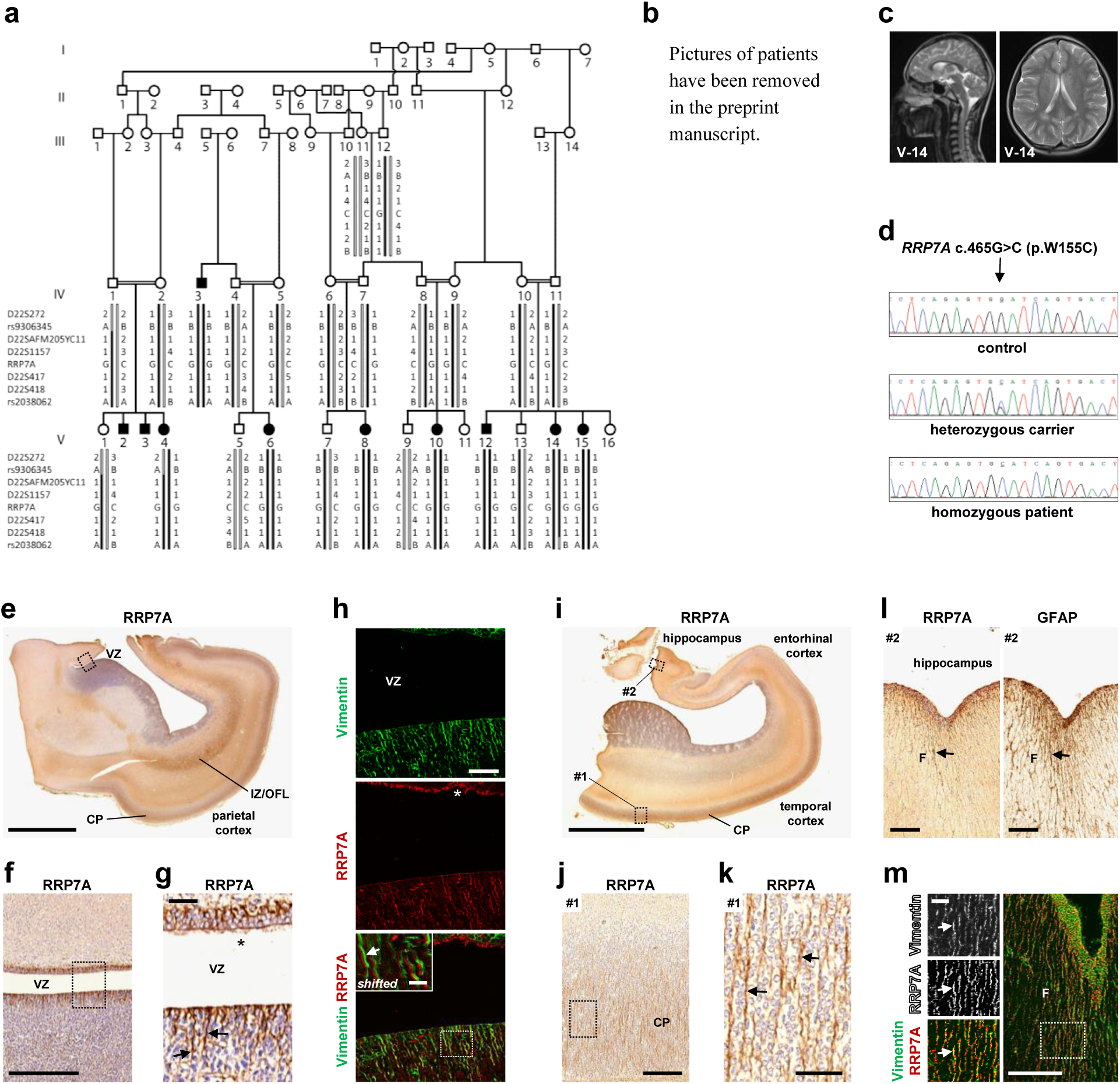
RRP7A is associated with MCPH and expressed in RGCs in the developing human neocortex. **a**, Five-generation pedigree of a consanguineous Pakistani family with ten affected individuals (black filled symbols). STR and single nucleotide polymorphism marker haplotypes of all analysed individuals are shown below each symbol. Diseased haplotype is marked as filled black bar. **b**, Clinical pictures of four affected family members presenting with microcephaly and sloping forehead. **c**, Cerebral MRI scans of patient V-14. **d**, DNA sequence of homozygous control, heterozygous carrier and homozygous patient with c.465G>C (p.W155C) mutation in exon 5 of *RRP7A*. **e-m**, RRP7A expression in RGCs and cilia of human fetal brains aged 19 wpc. **e**,**f**, DAB staining depicting the expression of RRP7A in the human parietal cortex in low (e) and higher (f) magnification of the ventricular zone (VZ). Scale bars, 5 mm (**e**) and 0.5 mm (f). **g**, Higher magnification of the zone boxed in **f**. RRP7A reactivity is high in RGCs (arrows) and cilia (asterisk). Scale bar, 40 µm. **h**, IFM analysis of the region depicted in (g) showing expression of RRP7A (*red*) in RGCs marked with Vimentin (*green*) and localization to cilia (asterisk). Scale bar, 50 µm. Insert: Shifted overlay of region boxed in the merged panel showing RRP7A expression in RGCs (arrow). Insert scale bar, 10 µm. **i**, DAB staining depicting the expression of RRP7A in RGCs in a section of the temporal cortex and hippocampal formation. Scale bar, 5 mm. **j**,**k**, Higher magnifications of the zone boxed #1 in (i) showing expression of RRP7A in RGCs at the cortical plate (CP). Scale bar, 0.2 mm. **k**, Higher magnification of the zone boxed in (j). Scale bar, 50 µm. **l**, Higher magnification of the zone boxed #2 in (i) showing expression of RRP7A (left panel) to RGCs (arrow) at the outer surface along the hippocampus (F: Fimbria). RGCs (arrow) in this region are further marked with Glial Fibrillary Acidic Protein (GFAP) (right panel). Scale bars, 0.2 mm. Low magnification with GFAP staining is also shown in Supplementary Fig. 2a. **m**, IFM analysis of the area shown in (l) showing expression of RRP7A (*red*) to RGCs (arrows) marked with Vimentin (*green*). Inserts scale bar, 20 µm. The three left panels represent the zone boxed in the right panel. Abbreviations: IZ/OFL: Intermediate zone/outer fibre layer.

To evaluate the spatial expression of RRP7A in the developing human brain, DAB staining (bright-field immunohistochemistry) and immunofluorescence microscopy (IFM) were performed on sections of midgestation human fetal brains, in which all proliferative zones of the developing cortex are represented^8^. Examination of the parietal cortex showed expression of RRP7A in RGCs at the VZ, ISVZ, OSVZ, intermediate zone/outer fibre layer, subplate and cortical plate as well as localization to ependymal cilia at the VZ (Fig. 1e-h). See Supplementary Fig. 2a for indications of the individual zones stained for Vimentin that marks RGCs. Further, evaluation of the temporal cortex and hippocampal formation revealed RRP7A localization to RGC fibres in the cortical plate (Fig. 1i-k) and at the outer surface along the fimbria (Fig. 1l,m), which is devoid of neurons at this stage as confirmed by lack of βIII-tubulin staining (Supplementary Fig. 2b). RRP7A also localizes to other brain areas and cell types, including the meninges and endothelial cells lining the brain blood vessels (Supplementary Fig. 2c). The identification of OSVZ as the key germinal zone responsible for neocortical expansion^9^ and the subsequent characterization of its population of Hopx-positive basal RGCs provided additional insight in cortical and evolutionary neocortical expansion^10^. The prominent RRP7A immunoreactivity of apical truncated RG and in particular bRG following the discontinuity of the radial glia scaffold in human neocortex present at midgestation^2^ indicates an important role in neural and neuronal development.

To assess the role of RRP7A in proliferation and neurogenesis, we used the CRISPR/Cas9 system to induce random deletions in exon 2 of the *Rrp7a* gene in P19CL6 cells, a mouse teratocarcinoma stem cell line that differentiates into neurons upon retinoic acid (RA) stimulation^11^ (Supplementary Fig. 3a,b). Analysis of 94 clones revealed high genome editing efficiency, with an *Rrp7a* alteration in 53% of cell clones (data not shown). However, none of the observed mutations caused frameshift in both copies of *Rrp7a*. We therefore chose two clones (P19CL6^*Rrp7a*Δ1/Δ18^ and P19CL6^*Rrp7a*Δ8/Δ33^), both of which contain two mutant *rrp7a* alleles (an in-frame and a frameshift deletion, respectively, (Supplementary Fig. 3c)), and analysed these in parallel with two wild-type (WT) clones (P19CL6^*Rrp7a*WT#1^ and P19CL6^*Rrp7a*WT#2^), which readily differentiate into neurons (Supplementary Fig. 3d). As expected, WB analysis showed that the mutant clones express reduced levels of lower molecular mass species of RRP7A compared to the WT clones (Fig. 2a) and IFM analysis showed that these clones proliferate slower than the WT clones (Fig. 2b,c). To investigate whether RRP7A is required for neuronal differentiation, WT and mutant clones were cultured to the same confluency of ca. 30% followed by RA stimulation (Fig. 2d). In contrast to P19CL6^*Rrp7a* WT#1^, which formed neurons at linings of cell clusters at day 6 of stimulation and produced elaborate neuronal networks positive for Microtubule Associated Protein 2 (MAP2) and βIII-tubulin at day 9, differentiation was prominently reduced in the mutant clones (Fig. 2d,e). Impaired neurogenesis in the mutant clones was confirmed by SDS-PAGE and WB analyses, showing that P19CL6^*Rrp7a*WT#1^ but not P19CL6^*Rrp7a*Δ8/Δ33^ displayed significant upregulation of the neuroectodermal lineage marker Paired Box 6 (PAX6) as well as βIII-tubulin and MAP2 during RA stimulation (Fig. 2f-h). Taken together, our data suggest that RRP7A promotes cell proliferation and is required for timely onset of neurogenesis and formation of elaborate networks of post-mitotic neurons.

**Fig. 2:**
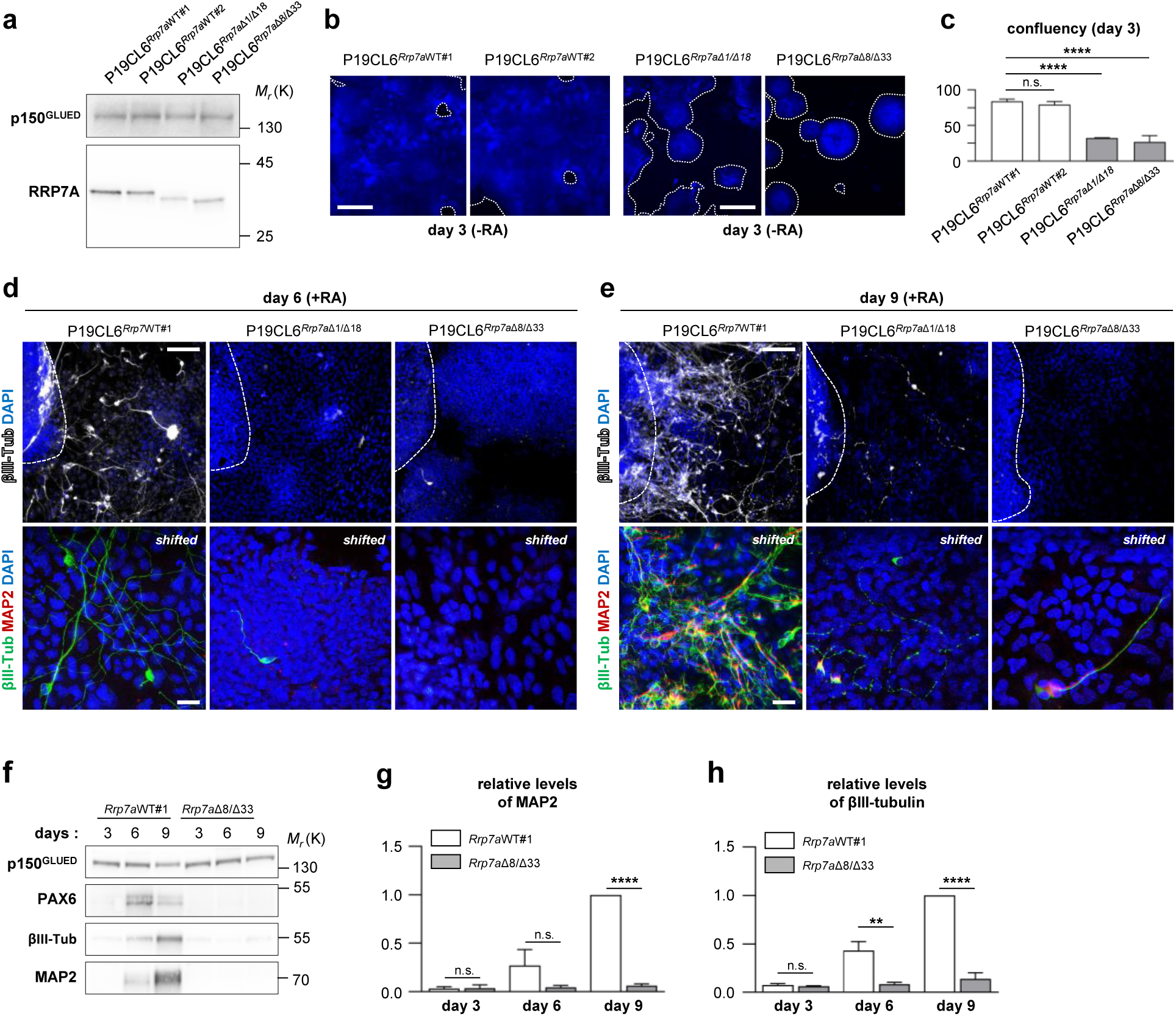
Mutation of RRP7A inhibits proliferation and neurogenesis. **a**, WB analysis of RRP7A expression in CRISPR clones of P19CL6 mouse stem cells holding WT RRP7A (P19CL6^*Rrp7a*WT#1^ and P19CL6^*Rrp7a*WT#2^) or mutated RRP7A (P19CL6^*Rrp7a*Δ1/Δ18^ and P19CL6^*Rrp7aA*Δ8/Δ33^). **b**, IFM analysis of CRISPR clone proliferation at day 3 post seeding in the absence of retinoic acid (RA). Nuclei were stained with DAPI (blue). Dotted lines outline cell colonies. Scale bars, 1 mm. **c**, Quantification of proliferation of CRISPR clones measured by cell coverage (confluency) (n=3). **d**,**e**, IFM analysis of the ability of CRISPR clones to undergo neurogenesis after 6 (d) and 9 (e) days of RA stimulation. The stippled lines in the upper panels indicate the lining of cellular clusters from where neurons are formed in WT clones. Neurons are marked with anti-ßIII-tubulin (upper panels: *white*; lower panels: *green*) and anti-MAP2 (lower panels: *red*), and nuclei are stained with DAPI (*blue*). Lower panels are shown as shifted overlays between ßIII-tubulin and MAP2. Scale bar upper panels, 0.1 mm and lower panels, 40 µm. **f**, WB analysis of WT clone P19CL6^*Rrp7a*WT#1^ and mutant clone P19CL6^*Rrp7a*Δ8/Δ33^ at day 3, 6, and 9 of RA stimulation; antibodies as indicated. **g**, Quantification of relative levels of MAP2 shown in (f) (n=3). **h**, Quantification of relative levels of ßIII-tubulin shown in (f) (n=3).

A comprehensive cell-based siRNA screen previously implicated RRP7A in ribosome assembly^12^ and the W155 residue is involved in the interaction of RRP7A with NOL6 (UTP22)^13^. Since defective ribosome biogenesis is associated with cell cycle defects as well as neurodevelopmental disorders in combination with additional abnormalities^14, 15^, we investigated if the p.W155C mutation in RRP7A affects rRNA processing using patient-derived (human) dermal fibroblasts (HDFs). Initially, we compared the expression of RRP7A in control and patient HDFs using WB and qRT-PCR analyses and observed reduced RRP7A protein levels, but normal mRNA levels, in the latter (Supplementary Fig. 4a,b). Addition of proteasome inhibitor MG-132 restored RRP7A levels in patient HDFs to normal (Supplementary Fig. 4a), suggesting that the p.W155C mutation leads to proteolytic degradation of RRP7A. IFM analysis of control HDFs, either cultured in the presence of serum or synchronized to growth arrest by serum depletion, showed that RRP7A is localized in the cytosol and predominantly to nucleoli, which is the principal site of ribosome biogenesis, as well as to the centrosome and primary cilium (Fig. 3a-c). Similar results were obtained for patient-derived HDFs except that the RRP7A signal in nucleoli was significantly reduced in these cells compared to controls (Fig. 3d-f). This phenotype was recapitulated in hTERT RPE-1 (RPE-1) cells expressing FLAG-RRP7A-GFP^WT^ or FLAG-RRP7A-GFP^W155C^ (Supplementary Fig. 4c-e), indicating that the p.W155C mutation impairs recruitment to nucleoli.

**Fig. 3:**
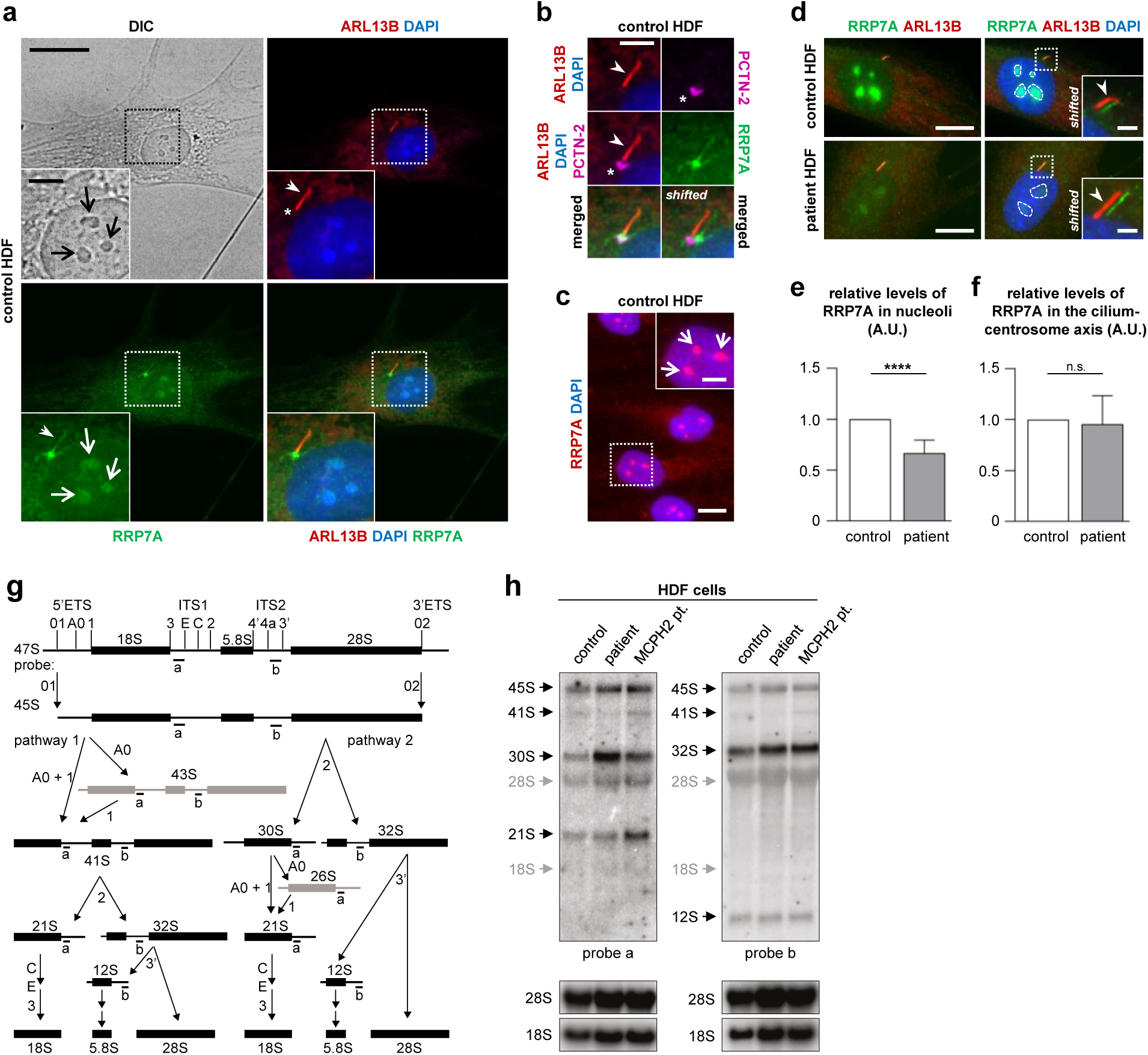
RRP7A localizes to nucleoli, centrosomes and primary cilia, and nucleolar localization and rRNA processing are disrupted in patient HDFs. **a**, IFM analysis on the localization of RRP7A (*green*) to nucleoli (open arrows, light microscopy (LM)), the primary cilium (closed arrows, ARL13B, *red*) in serum-depleted cultures of control human dermal fibroblasts (HDF). The nucleus (nu) is stained with DAPI (*blue*). Scale bar, 20 µm. Insert scale bar, Scale bar, 5 µm. **b**, Zoomed image of the cilium-centrosome axis depicted in (a) with the ciliary base/centrosome marked with anti-Pericentrin-2 (asterisk, PCTN-2, *magenta*). Scale bar, 5 µm. **c**, IFM analysis of localization of RRP7A (*red*) to nucleoli (open arrows) in serum supplemented cultures. Scale bar, 20 µm. Scale bar in zoomed image, 5 µm. **d**, IFM analysis on the localization of RRP7A (*green*) to nucleoli (stippled-lined areas) and the primary cilium (ARL13B, *red*) in serum-depleted cultures of control HDFs (left panel) and patient HDFs (right panel). Scale bars, 10 µm. The insert shows magnification of boxed areas with shifted overlays between RRP7A (*green* closed arrow) and ARL13B (*red* closed arrow). The nucleus (nu) is stained with DAPI. Scale bars, 2 µm. **e**, Quantification of the relative levels of RRP7A in nucleoli in control and patient HDFs shown in (d) (n=5). **f**, Quantification of the relative levels of RRP7A in nucleoli at the cilium-centrosome axis in control and patient HDFs shown in (d) (n=4). **g**, Map of the human pre-rRNA transcript with annotated processing sites and a simplified outline of the two main processing pathways with short-lived precursors in grey. In pathway 1, cleavage of 45S occurs at sites A0 and 1 to produce 41S that is subsequently split into two parts by cleavage at site 2 to yield the direct precursors for the small and large subunit rRNAs, respectively. In pathway 2, this order is reversed, such that 45S is first cleaved at site 2 yielding precursors for the small and large subunit rRNAs. The target sites of oligo nucleotide probes a and b are indicated on the map. **h**, Northern blots of parallel gel runs of total RNA samples from control and patient HDFs with mutations in RRP7A (patient) and WDR62 (MCPH2 pt.), respectively. Black arrows indicate processing intermediates inferred from the analyses and grey arrows mark the migration of mature rRNA species as inferred from re-probing of the filters with probes targeting these RNA species.

To investigate rRNA processing, total RNA from control and patient HDFs was analysed by northern blotting using oligo nucleotide probes targeting internal transcribed spacers 1 (ITS1) and 2 (ITS2). Fibroblasts from an MCPH patient with a mutation in the *WDR62* gene^16-18^, which is unrelated to ribosome biogenesis, was included as a control. In human pre-rRNA processing, the 47S precursor is first cleaved at sites 01 and 02 to form 45S, followed by processing according to one of two main pathways (Fig. 3g)^19-21^. The processing pattern in the *RRP7A* mutant differs from WT and the *WDR62* mutant in three ways (Fig. 3h). First, there is a depletion of 41S in pathway 1 (probes a and b). Second, there is an accumulation of 30S in pathway 2 (probe a). Third, there is a depletion of 21S in both pathways (probes a and b). These observations are consistent with a deficiency in cleavage at site A0 in formation of 18S rRNA. Accumulation of 45S or a shift towards increased use of pathway 2 could not be resolved due to co-migration of 47S and 45S. In contrast to the deficiencies in the pathway leading to 18S rRNA, the pathway leading to 28S appeared unaffected as evidenced by equal signals corresponding to 32S and 12S, respectively. In siRNA knockdowns of RRP7A in HeLa cells, primary lung fibroblasts, and HTC-116 colon carcinoma cells, the phenotype was characterized as “21S down, 18S-E down”^12^. Furthermore, re-inspection of the published northern blots^12^ suggest depletion of 41S and accumulation of 30S in RPP7A depleted cells. Thus, we conclude that the *RRP7A* mutation results in defective rRNA processing due to a deficiency in cleavage at site A0.

Since *Rrp7a* mutant P19CL6 clones and *RRP7A* patient HDFs generally proliferated slower than controls (data not shown) we speculated whether the observed defect in ribosome biogenesis in *RRP7A* mutant cells is correlated with a delay in cell cycle progression. To test this we monitored the assembly and resorption of the primary cilium, which is formed during quiescence to coordinate developmental signaling^22^ as well as to retain cells in G_0_/G_1_^23, 24^. Aberrant regulation of cilium resorption is thus linked to defects in proliferation-differentiation decisions of RGCs in the developing mouse neocortex^24^, and MCPH patient fibroblasts carrying a mutation in *CENPJ* present excessively long cilia, resulting in delayed ciliary disassembly and S phase entry, which in NPCs leads to premature differentiation and reduced brain size^25^. In cultured fibroblasts, cilium formation and resorption can be induced by serum withdrawal and re-addition, respectively. Following initial serum-induced cilium disassembly at G_0_/G_1_, cells are transiently reciliated to be deciliated again during S/G_2_ concomitant with DNA synthesis^26^. Interestingly, IFM analysis showed that RRP7A patient HDFs grown in serum enriched medium, which normally suppresses ciliogenesis and promotes cilium resorption, display significantly higher ciliation frequency than controls (time=0 h; Fig. 4a,b). In contrast, serum-depleted cells showed similar frequency and length of primary cilia in control and patient HDFs (Fig. 4b,c), suggesting that the RRP7A mutant HDFs fail to deciliate during conditions permissive for proliferation. To address ciliary resorption, cells were deprived of serum for 48 hours and then grown 24 hours in serum-enriched medium. In accordance with previous studies^26^, ciliary resorption in control HDFs proceeded in two waves, first ciliary length was reduced after 6 hours of serum stimulation, followed by transient elongation at 12 hours and complete ciliary resorption at 24 hours post serum re-introduction (Fig. 4d-f). Patient HDFs also displayed shortened cilia at 6 hours of serum stimulation, but cilia proceeded to elongate at 12-24 hours post serum-introduction, which was associated with a high frequency of ciliated cells even after 24 hours (Fig. 4d-f). WB analysis revealed that patient cells during serum re-introduction display delayed phosphorylation of retinoblastoma protein (RB), which marks cell cycle progression (Fig. 4g,h). These results support the conclusion that the p.W155C mutation in RRP7A compromises cell cycle progression, which is coupled to the second wave of ciliary resorption. Our combined results from patient HDFs and P19CL6 cells thus indicate that RRP7A invalidation leads to a reduction in both proliferation and neuronal differentiation.

**Fig. 4:**
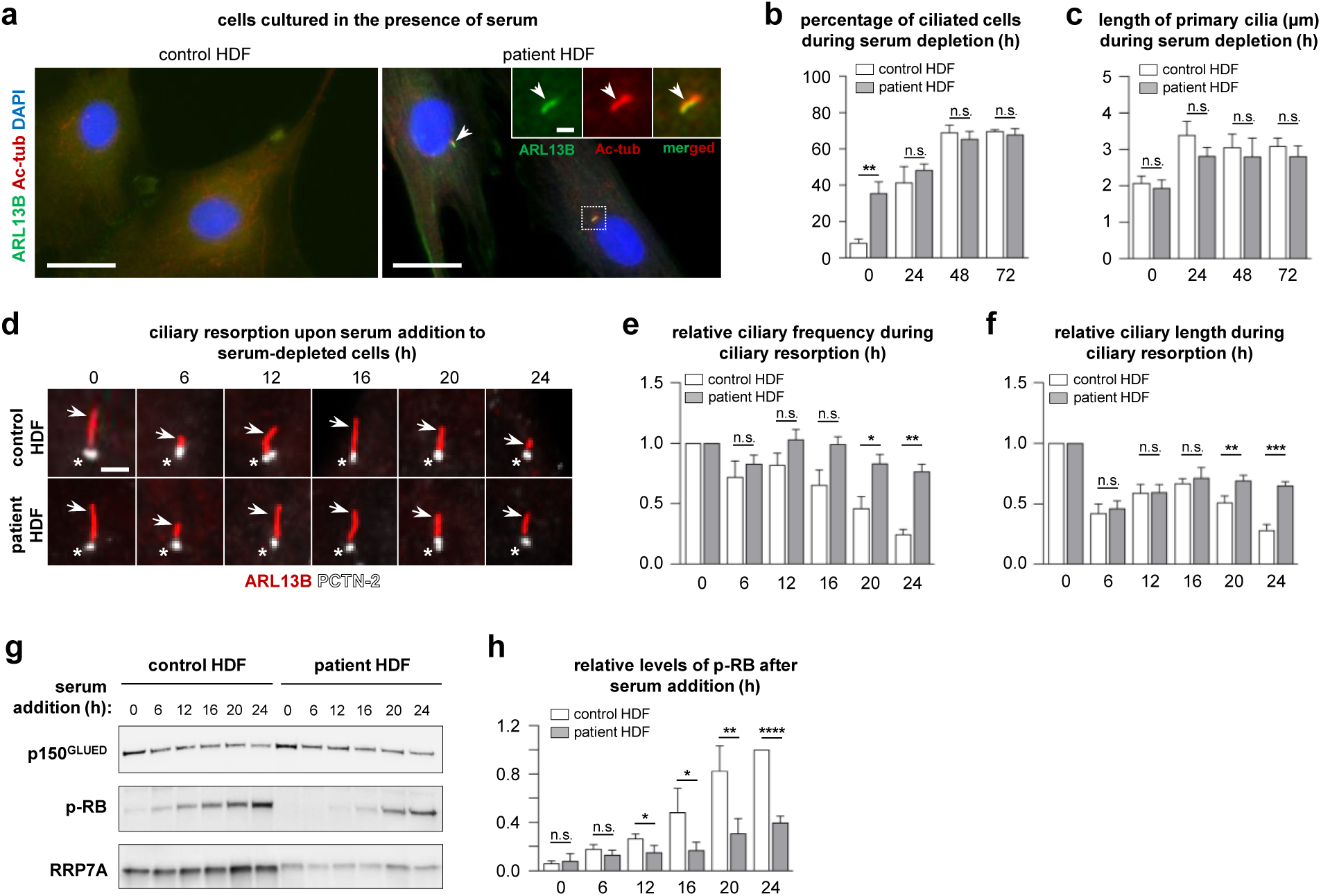
Patient HDFs display defects in resorption of primary cilia and cell cycle entry. **a**, IFM analysis of primary cilia in control and patient dermal fibroblasts (HDFs) cultured in the presence of serum. Primary cilia (arrows) are marked with anti-ARL13B (*green*) and anti-acetylated α-tubulin (Ac-tub, *red*), and nuclei are stained with DAPI (*blue*). Scale bars, 40 µm. Scale bar in zoomed image, 2 µm. **b**, Quantification of ciliary frequency in control and patient HDFs serum depleted for 0-72 hours (n=3). **c**, Quantification of ciliary length in control and patient HDFs serum depleted for 0-72 hours (n=4). **d**, Overview of ciliary lengths in control HDFs (upper panels) and patient HDFs (lower panels) subjected to serum depleted for 48 hours followed by serum addition for 0-24 hours. Cilia are marked with anti-ARL13B (arrows, *red*), centrosomes (ciliary basal bodies) are marked with anti-PCTN-2 (asterisks, *white*) and nuclei are stained with DAPI (*blue*). Scale bar, 2 µm. **e**, Quantification of cilia frequency based on data presented in (d) (n=3). **f**, Quantification of cilia length based on data presented in (d) (n=3). **g**, WB analysis of the level of retinoblastoma protein phosphorylation (p-RB) and RRP7A expression in control and patient HDF cells subjected to serum depleted for 48 hours followed by serum addition for 0-24 hours. **h**, Quantification of relative levels of p-RB shown in (g) (n=4).

Finally, to investigate the role of RRP7A in brain development *in vivo*, we used zebrafish as a model. Whole mount *in situ* hybridization analysis of 2 days post fertilization (dpf) zebrafish embryos showed *rrp7a* expressed in the head of the embryo, including the brain and the eyes (Fig. 5a). We proceeded to study developmental defects associated with *rrp7a* depletion using a zebrafish line carrying a frameshift mutation in *rrp7a* (Supplementary Fig. 5a,c). Homozygous mutant progeny presented with a phenotype resembling primary microcephaly, characterized by significantly reduced size of the brain, but otherwise no gross abnormalities (Fig. 5b-f). In line with the observed defects in neurogenesis and cell cycle control in mutant P19CL6 cells and HDFs, reduced brain size was accompanied by a reduction in the number of cells in the pallium of 3 dpf *rrp7a*^*-/-*^ mutants embryos (Fig. 5g) as well as a marked reduction in the expression of the neural progenitor marker *neurod1* and proliferation marker *pcna* in the developing brains of *rrp7a*^*-/-*^ mutants (Fig. 5h). Ultimately, to consolidate the observations from patient HDFs, we analysed rRNA processing in WT and *rrp7a*^-/-^ zebrafish. The embryos were sorted according to phenotype based on morphological criteria and RT-PCR confirmed aberrant splicing of *rrp7a* mRNA in embryos with mutant phenotype (Supplementary Fig. 5b). Northern blotting analysis showed an altered pattern of processing intermediates and a pronounced depletion of mature 18S rRNA in the knock-out compared to WT fish while mature 28S and 28S processing intermediates appeared unaffected (Fig. 5i).

**Fig. 5:**
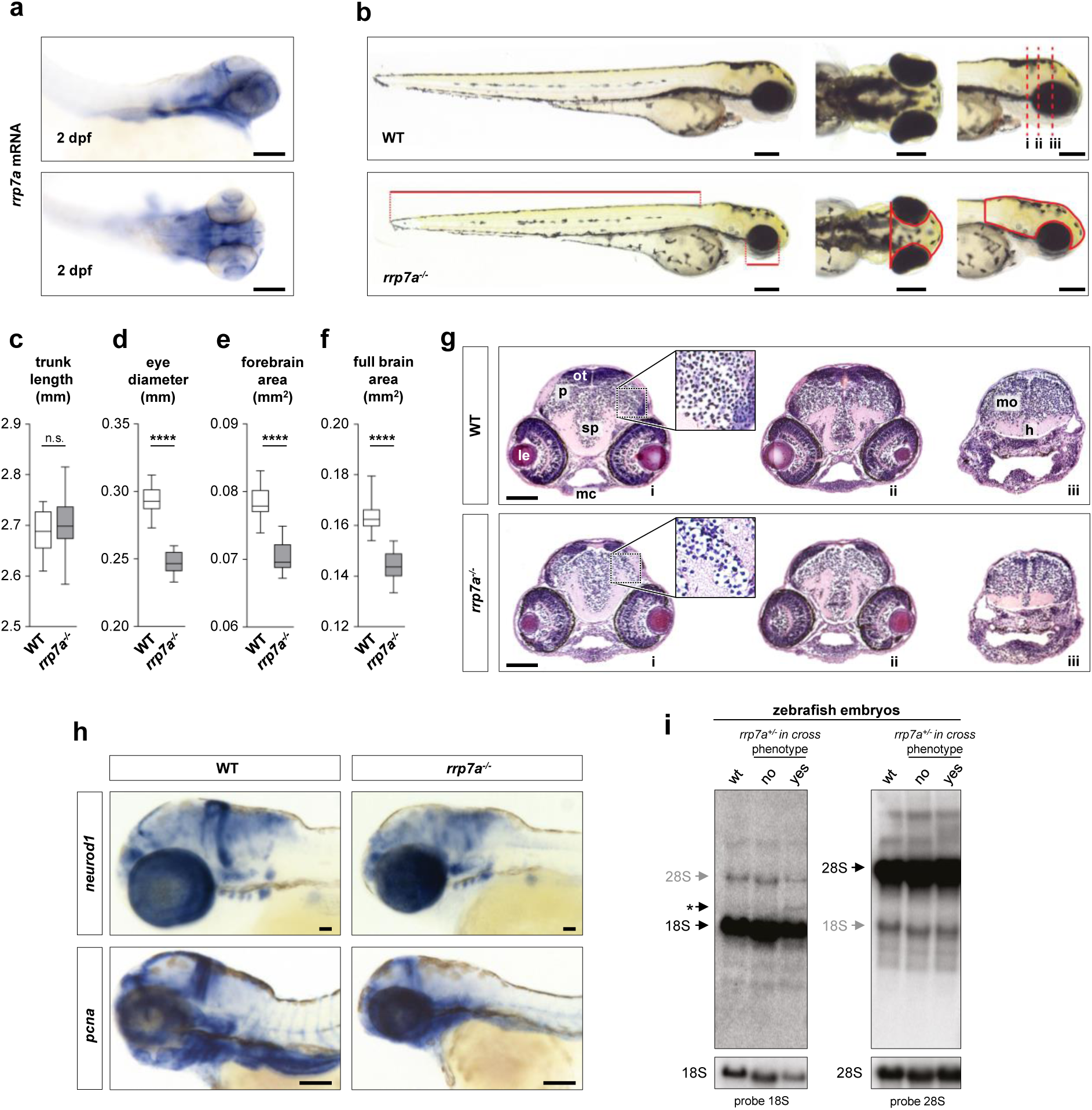
Knockout of Rrp7a in zebrafish causes brain defects resembling primary microcephaly. **a**, Expression of *rrp7a* in 2 dpf zebrafish embryos by *in situ* hybridization. Scale bars, 0.1 mm. **b**, Bright field images showing the morphology of 3 dpf wild-type (WT) and homozygous *rrp7a*^*-/-*^ mutant zebrafish embryos. Scale bars, 0.2 mm. (**c-f**) Quantification of morphological differences between WT and mutants. Trunk length **c**, eye diameter **d**, forebrain area **e** and full brain area **f** were quantified by image analysis. Measured regions are indicated in **b** (red solid lines). **g**, Hematoxylin and eosin (HE) staining of transverse tissue sections from a 3 dpf WT (upper panels; i-iii) and *rrp7a*^*-/-*^ mutant (lower panels; i-iii) zebrafish embryo. Location of sections is indicated in (b) (red dotted lines). Note the reduction in cells in the pallium (p) of mutants compared to WT (marked in zoomed box). Abbreviations: h: hypothalamus, le: lens, mc: Meckel’s cartilage, mo: medulla oblongata, ot: optic tectum, sp: subpallum. Scale bars, 0.1 mm. **h**, *in situ* hybridization analysis of *neurod1* and *pcna* mRNA expression in WT and *rrp7a*^*-/-*^ mutants at 3 dpf. Scale bars, 0.1 mm. **i**, Northern blot of total RNA from pooled zebrafish embryos analysed with 18S rRNA and 28S rRNA probes, respectively. The asterisk marks a band accumulating in *rrp7a*^*-/-*^ embryos only. The panels at the bottom of the figure were adjusted in exposure to highlight the depletion of 18S rRNA relative to 28S rRNA in *rrp7a*^*-/-*^ embryos.

In conclusion, we have identified *RRP7A* as a novel disease gene in MCPH. Analyses of human fetal brains, zebrafish, cell models, and primary cells from patients showed that RRP7A regulates brain development, neurogenesis, ciliary resorption and cell proliferation and provide evidence for defective ribosome biogenesis as pathophysiological mechanism. While RRP7A was observed to localize to cilia and centrosomes in addition to nucleoli, it is currently unclear if and how the pool of RRP7A at the cilium-centrosome axis impinges on the disease phenotype. It will be interesting to address this in future studies; in particular whether cilium-localized RRP7A might directly promote ciliary disassembly prior to mitosis and/or regulate developmental signalling associated with RGC/NPC function and neurogenesis.

## Acknowledgements

We thank Maria S. Holm, Lillian E. Rasmussen, Pernille S. Froh, Troels Askhøj Andersen and Martin Juel Vilhelm for excellent technical assistance. This study was supported by grants from the Lundbeck Foundation (R221-2016-922), the University of Copenhagen Excellence Programme for Interdisciplinary Research, the Novo Nordisk Foundation (NNF14OC0011535 and NNF15OC0016886), Independent Research Fund Denmark (DFF-6108-00457), and the Carlsberg Foundation (CF15-0425 and CF18-0532) (to S.T.C., L.A.L. and L.B.P.); grants from the Lundbeck Foundation (R198-2015-174) and the Danish Cancer Society (R167-A10943-17-S2) (to H.N. and N.K.); a grant from the Independent Research Fund Denmark (DNRF107) (to LH); a grant from The Lundbeck Foundation (R209-2015-2604) (to C.D.).

## Author Contributions

L.A.L., S.T.C., M.F. and L.L. conceived the study. S.T.C. and L.A.L. directed the experiments and wrote the manuscript with input from all authors. All authors provided intellectual input to the project. M.F. and S.M.B. identified and performed clinical investigations of the family. M.F., Y.M., H.E., A.F and L.H. performed genetic analyses. L.L. and K.M. performed immunohistochemistry on fetal human brain sections, which were provided by K.M. L.L., C.K and V.S.N. constructed and analysed P19CL6 mutant lines. L.L. and M.F. analysed localization and expression of RRP7A in HDF and RPE-1 cell lines. H.N. and N.K. analysed pre-rRNA processing in HDFs and zebrafish. L.L. and C.K. analysed ciliary formation, ciliary resorption and cell cycle progression in HDFs. C.D., M.M. and S.S. performed zebrafish experiments.

## Materials and Methods

### Patient samples

The study followed the declaration of Helsinki and was approved by institutional ethical committees. The five generation family originates from the rural area in the Rahim Yar Khan district, in Punjab province of Pakistan. Informed consent was obtained from all participating individuals or their parents for the collection of blood and/or skin biopsy samples, genetic analyses, and publication of photos and genetic information. Blood samples from all available family members were collected in separated appropriate tubes for genomic DNA/RNA isolation. Skin biopsies from affected individuals were collected in DMEM for cell cultures. Genomic DNA was isolated from peripheral blood cells using phenol-chloroform extraction.

### Magnetic resonance imaging

A magnetic resonance imaging (MRI) scan was performed for one affected individual at the Sheikh Zayed Hospital, Rahim Yar Khan. Multiplanar, multisequential FLAIR and diffusion images of brain MRI were obtained in axial, coronal and sagittal views and Images were analysed using *syngo* fastView (version 1.0.0.3, Siemens AG).

### Genome-wide SNP array mapping and linkage analysis

To identify regions of homozygosity, deletions, or duplications, samples were genotyped on Genome-Wide Human SNP array 6.0 (Affymetrix Inc., Santa Clara, CA, USA). The analysis was performed in four patients (one from each loop) according to the manufacturer’s standard protocol (Affymetrix). Data analysis was performed using Chromosomal analysis suite (ChAS) and FASTLINK software tools (Schaffer et al 1994). Polymorphic STR markers (D22S272, D22SAFM205YC11, D22S1157, D22S417, D22S418, primers are available upon request) were used to genotype all available family members to identify and map the linked chromosomal region. STR markers were PCR amplified and alleles were separated in an ABI 3130*xl* Genetic Analyzer (Applied Biosystems, Foster City, CA, USA) according to the manufacturer’s instructions. Results were analysed using Peak Scanner Software, version 1.0 (Thermo Scientific, Waltham, MA USA).

### Genetic analysis

Direct sequencing of *ASPM* and linkage analysis of *WDR62* and *CENPJ*, using microsatellite markers flanking each gene, was used to exclude the most common MCPH genes in the Pakistani population. Following exclusion of the most commonly involved genes, linkage analysis was performed using Genome-Wide Human SNP 6.0 arrays. This analysis excluded linkage of rare copy number variants (CNVs).

Whole exome sequencing (WES) was performed on genomic DNA extracted from an affected individual. Capture was performed using the Nextera Rapid Capture Exome Enrichment Kit and sequencing was done using a paired-end 2 x 100 bp protocol using an Illumina Hi-seq 2000 system (Illumina, San Diego, CA, USA). Data was filtered and aligned to the Human Genome reference (hg38) using Burrows Wheel Alignment. Variants were analysed using the Genome Analysis Toolkit. Variants with minor allele frequency of <0.01 were filtered against known databases of genomic variants (dbSNP138, ExAC). The *RRP7A* mutation c.465G>C was confirmed by direct Sanger sequencing of a PCR amplified exon-fragment in all available samples from patients and unaffected family members. The following primers were used: 5’-GACTACATGTGTCACCCAGC-3’ (forward) and 5’-GGATCTCCTTAGCACCTGTG-3’ (reverse). Sequencing was performed in an ABI 3130*xl* genetic analyzer using the ABI Prism Big Dye Terminator v3.1 Cycle Sequencing Kit (Applied Biosystems) according to the manufacturer’s protocol. Sequences were analysed using chromaspro software (chromaspro 2.1, Technelysium Pty Ltd, South Brisbane, Australia). The c.465G>C mutation leads to the introduction of an *Hpy*CH4V restriction enzyme site in the mutant. Analysis of the mutation in DNA samples obtained from 300 ethnically matched Pakistani control individuals were performed by restriction digestion on amplified PCR products, using the HpyCH4V restriction enzyme. HpyCH4V restriction enzyme digestion products were analysed on 1.5% agarose gels.

### Multiple alignment and homology modelling of RRP7A

Multiple protein sequences alignments to determine the conservation of the mutated amino acid (p.W155C) was done using the online alignment tool (http://www.uniprot.org/). RRP7A (>NP_056518.2) has 23% sequence similarity with the Ribosomal RNA-processing protein 7 [(RRP7) *S. cerevisiae*].

### Total RNA isolation, RT-PCR and qRT-PCR

To check for RRP7A splicing errors, total RNA was isolated from whole blood obtained from a patient (V.14) and a healthy control individual. Blood for RNA was collected in PAXgene blood RNA tubes, and was isolated using a PAXgene^®^ Blood RNA Kit (PreAnalyticX). One µg of total RNA was reverse transcribed using SuperScript™ II (Invitrogen, Carlsbad, CA), and oligo-dT primers (Invitrogen). To detect aberrant splicing, full length *RRP7A* gene products were PCR-amplified from first strand cDNA with gene specific primers 5’-ATGGTGGCGCGCAGGAGGAA-3’ (forward) and 5’-GTACGGTCGGAATTTGCG-3’ (reverse). The PCR products were analysed on 1% agarose gels, and subsequently directly sequenced in both directions. To check for the analysis of ribosomal RNA processing defects, total RNA was extracted from fresh cultures of primary fibroblasts obtained from an affected individual (V.14), an MCPH patient homozygous for *WDR62* mutation (loss of function mutation due to microduplication (unpublished data)) and a normal control using QIAzol^®^ lysis reagent (Qiagen Inc., Valencia, CA) in accordance with the manufacturer’s guidelines. Quality of extracted RNAs were checked by running samples in 1% agarose gels and quantification was done using NanoDrop™ ND-1000 spectrophotometer (Thermo Fisher Scientific Inc., Wilmington, USA). In order to assess expression of *RRP7A* mRNA in primary fibroblast cultures, qRT-PCR was performed using Brilliant III Ultra-Fast SYBR® Green QPCR master mix (Agilent Technologies). The qRT-PCR reactions were performed in triplicate and were analysed on an ABI 7500 fast system. The data was normalized against the average expression value of two housekeeping genes (*GAPDH* and *B2M*, primer sequences available upon request).

### Human brain tissue samples and immunohistochemistry

Eight brains from 7 weeks post conception (wpc) to 21 wpc were analysed. A 19 wpc midgestation fetus was chosen for the entire immunohistochemical analysis as material from all brain regions was available and all cell types present at this stage. The material was obtained from legal abortions. Written and oral informed consent was given according to the Helsinki declaration II, approved by the Research Ethics Committee of the Capital Region (KF–V.100.1735/90). Immediately following operation, the samples were dissected into blocks and fixed for 12–24 hours at 4°C in either 10% neutral buffered formalin, 4% Formol-Calcium, Lillie’s or Bouin’s fixatives. The specimens were dehydrated with graded alcohols, cleared in xylene, and paraffin embedded. Serial sections, 3–10 µm thick, were cut in transverse, sagittal, or horizontal planes, placed on salinized glass slides, and used for immunohistochemical experiments^27^. Immunohistochemical analyses were performed on deparaffinised sections of human brain 19 wpc. Samples were carefully treated in heat-induced antigen retrieval using citrate buffer pH 6.0, prior to blocking of unspecific binding and incubation with primary antibodies overnight, according to previously described protocols^27^. Immunofluorescence staining was prepared as for bright field microscopy. Primary antibodies were mixed in proper dilutions, and tissue was incubated over-night. Subsequently, sections were washed in TBS and incubated for 45 minutes at RT in Alexa Flour antibodies of the desired wavelengths. Antibody dilutions and manufacturers are listed in Supplementary Tables 2 and 3. Sections were mounted using Shandon™ ImmuMount (Thermo Fisher Scientific).

### Cell cultures for *in vitro* neurogenesis analysis

P19CL6 murine embryonal carcinoma stem cells (Murofushi, Kimiko Japan, ref. 2406 3467) were cultured at 37°C and 5% CO_2_ in MEMalpha medium (Invitrogen 22561-05, Carlsbad, CA, USA) supplemented with 10% heat inactivated fetal bovine serum (FBS, Gibco) and 5% penicillin/streptomycin (p/s) (Gibco), and seeded at 1000 cells/cm^2^. Cell density measurements and induction of neuronal differentiation with 1 mM retinoic acid (RA) (Sigma) were performed after 3 days of culturing in order to allow potential difference in cell proliferation to be evident. Cells were sampled for immunofluorescence microscopy (IFM) and western blotting (WB) analyses on day 0, 3, 6 and 9. Cells were counted using the NucleoCounter® NC-200™ automated cell counter from ChemoMetec, and days of differentiation were counted from the first day of RA treatment.

### Isolation and culturing of patient fibroblasts and hTERT RPE-1 cells

Skin punch biopsies were obtained from an affected individual homozygous for *RRP7A* c.465G>C and from a MCPH patient homozygous for mutation in *WDR62* mutation. The samples were treated with Dispase II (1.5 U/ml, Roche) in phosphate buffered saline (PBS) at 4°C for 3 hrs to separate epidermis from dermis. The dermis was then incubated at 37°C and 5% CO_2_ in Dulbecco’s modified Eagle’s medium (DMEM) (Invitrogen) supplemented with 10% FBS, 1% GlutaMAX™ (GIBCO^®^), 100 IU/ml penicillin, and 100 μg/ml streptomycin. Culture medium was changed after every 3-4 days and a pool of fibroblasts was expanded by further sub-culturing. Similarly, control fibroblasts were obtained using skin biopsy from 34 years old healthy control female of Asian origin. hTERT RPE-1 (RPE-1) cells were cultured in DMEM medium supplemented with 10% FBS and 5% p/s, medium was changed every 2-3 days.

### Generation of CRISPR clones of P19CL6 cells

CRISPR was performed as previously described^28^. In brief, 23 nt oligos targeting the second exon of *Rrp7a* were cloned into a pSpCas9(BB)-2A-GFP (PX-458) vector, amplified in *Eschericia coli* DH10α, and purified using NucleoBond Xtra Midi EF Kit (Machery-Nagel). Plasmid inserts were sequenced at Eurofins (MWG Operon, Ebersberg, Gernmany). P19CL6 cells were transfected using the Nucleofector device II (Amaxa, Biosystems Lonza, Switzerland) and Nucleofector Kit V (Amaxa Biosystems Lonza, Switzerland) as previously described^29^. Clones were evaluated by Indel Detection by Amplicon Analysis (IDAA) in which amplicons were labelled and DNA fragments were analysed by automated capillary electrophoresis for detection of indels induced by CRISPR/Cas9 mediated gene targeting of Rrp7a, as described^30^. In brief, tagged forward primer (5’AGCTGACCGGCAGCAAAATTGCCCGTGGATGGGTGGATG-3’) was used for amplification of the DNA fragment covering the sgRNA binding sites, and analysed using the ABI 3010 sequenator (ABI/Life Technologies, USA) under conditions specified by the manufacturer. Raw data was analysed using the Peak Scanner Software V1.0 (ABI/Life Technologies, USA). Clones were sequenced by Sanger sequencing using forward primer 5’-CCCGTGGATGGGTGGATG-3’ and reverse 5’-CACCAGGATTCTGTAGGGC-3’. Two guides were used to generate clone P19CL6^*Rrp7a*Δ8/Δ33^: Sg1 5’-AGACAGCACCGAGTTCGCCAAGG-3’ and Sg2 5’-CCCGATCGAACCCTTTTTATCCT-3’. Only Sg2 were used for clone P19CL6^*Rrp7a*Δ1/Δ18^. Two wild type clones (P19CL6^*Rrp7a* WT#1^ and P19CL6^*Rrp7a* WT#2^) were chosen as control cell lines, and their ability to differentiate was confirmed upon 9 days of RA treatment, in which both cell lines readily form elaborate networks of βIII-tubulin positive neurons (Supplementary Fig. 3e).

### Constructs

Using first strand cDNA obtained from a healthy donor as templates, full length coding sequence of the *RRP7A* (GenBank accession number NM_015703.4) was PCR amplified and cloned into the mammalian expression vector pEGFP-n1 using *Eco*RI and *Bam*HI restriction sites. Two full length wild type expression constructs for RRP7A were generated; one containing a Green Fluorescent Protein (GFP)-tag at the *C*-terminus and another containing a FLAG-tag at the *N*-terminus and a *C*-terminal GFP-tag, using pEGFP plasmid (Addgene). The mutation p.W155C was introduced into these vectors using QuikChange II Site-Directed Mutagenesis Kit (Agilent technologies Inc.). The primers used were 5’-GGCATTCACAAGTGCATCAGTGACTACGC-3’ (sense) and 5’-GCGTAGTCACTGATGCACTTGTGAATGCC-3’ (anti-sense). Expression constructs were verified by Sanger sequencing, and expressed in RPE-1 cells. Cells were seeded in DMEM without p/s, and transfected using FuGENE HD Transfection Reagent (Promega) when they reached approximately 60% confluence.

### SDS-PAGE and western blotting analyses

After washing in 1X ice cold PBS, cells were lysed in 1X SDS lysis buffer with Complete, EDTA-free Protease Inhibitor Cocktail (04693132001, Roche, Basel, Switzerland), with/without 2 mM sodium orthovanadate (S6508, Sigma). Cells were sonicated thrice for 5 seconds on ice (30 seconds for P19CL6 clones) and centrifuged at 20,000*g* for 20 min at 4°C (10°C for P19CL6 clones). Cleared supernatants were collected, and protein quantification was done using DC™ protein assay kit (Bio-Rad laboratories Inc. Hercules, CA, USA) according to the manufacturer’s instructions. Then, Samples were denatured in NuPAGE LDS sample buffer (NP0007, Invitrogen) at 95°C for 5 min. Then separated on gels (NuPAGE, Invitrogen) under reducing conditions and blotted onto nitrocellulose membrane (LC2006, Invitrogen). Blocking was done using 5% skim milk or bovine serum albumin (BSA) in TBST buffer. Membranes were incubated with horseradish peroxidase-conjugated secondary antibodies, developed using FUSION FX chemiluminescence system (Vilber Lourmat), and processed for presentations using Adobe photoshop CS6 version 13.0. Band intensities were analysed using ImageJ version 2.0 and statistical calculations were performed on n=3 or more. Details on antibodies concentrations and manufacturers are listed in Supplementary Tables 2 (primary antibodies) and 3 (secondary antibodies).

### Immunocytochemistry

IFM was performed as previously described^31^. In brief, cells were fixed in 4% PFA and permeabilized in 0.2% Triton X-100. Blocking was performed using 5% BSA, and cells were incubated in primary antibodies for 1.5 hours at RT or overnight at 4°C. Secondary antibodies were incubated for 45 min, and DAPI staining performed for 30 seconds prior to mounting in mounting media containing 2% N-propyl gallate and sealed with nail polish. Details on antibodies concentrations and manufacturers are listed in Supplementary Tables 2 and 3. Fluorescence and differential interference contrast (DIC) images were captured on a fully motorized Olympus BX63 upright microscope with an Olympus DP72 color, 12.8-megapixel, 4.140 x 3.096-resolution camera and with a fully motorized and automated Olympus IX83 inverted microscope with a Hamamatsu ORCA-Flash 4.0 camera (C11440-22CU). The software used was Olympus CellSens dimension, which was able to do deconvolution on captured z stacks, and images were processed for publication using Image J version 2.0 and Adobe Photoshop CS6. Quantifications of immunofluorescence intensities and cell confluency were performed in Cell Sense (Olympus). Careful outline of the cilium-centrosome axis as well as nucleoli were drawn using the outline tool, and mean intensities were obtained. ARL13B and Acetylated α-tubulin (Ac-tub) were used as ciliary markers to study the formation, length and disassembly of primary cilia. All data were gathered for n≥3.

### Northern blotting analysis of ribosomal RNA

Total RNA (7.5 µg) from human fibroblasts or zebrafish embryos was separated by denaturing gel electrophoresis on a formaldehyde denaturing 1% agarose gel and subsequently transferred to a positive charged nylon membrane (BrightStar-Plus, Ambion) by capillary blotting, followed by UV cross-linking. The probes (10 pmol each) were labelled with [γ-^32^-P]-ATP using T4 polynucleotide kinase (Thermo Fisher Scientific) and hybridized to the membrane in hybridization buffer (4X Denhardts solution, 6X SSC, 0.1% SDS), at 10°C below Tm of the probe for 16 hr. The membrane was then washed four times in washing buffer (3X SSC, 0.1% SDS) and exposed to a Phosphor Imager Screen, scanned by a Typhoon scanner (GE Healthcare) and analysed by ImageQuant software. The following probes were used for northern blotting analysis of human rRNA, probe a; 5’TGGGTGTGCGGAGGGAAGC targeting rRNA between processing site 3 and E in ITS1, probe b; 5’ACGCCGCCGGGTCTGCGCTTA targeting rRNA between processing site 4a and 3’ in ITS2, 5’CCAGACAAATCGCTCCACCAACTAAG, and 5’GCTCCCGTCCACTCTCGAC targeting 18S and 28S rRNA, respectively. The following two probes 5’CCAGACAAATCGCTCCACCAACTAAG, and 5’GCTCCCGTCCACTCTCGAC were used for northern blotting analysis of zebrafish 18S and 28S rRNA, respectively.

### Zebrafish maintenance

Embryos were maintained and staged as previously described^32, 33^ and all experiments were conducted according to the guidelines of the Danish Animal Experiments Inspectorate. The *rrp7a* (ENSDARG00000098934) mutant line (sa11429) was acquired from Zebrafish International Resource Center (ZIRC)^34^ F3 embryos were raised to adulthood and outcrossed to TL wild-type strain. All analyses were performed on F5 embryos, generated by crosses of heterozygous F4 adult fish.

### Zebrafish measurements and sectioning of embryos

Lateral and dorsal images of 3 days post fertilization (dpf) zebrafish embryos were taken under Zeiss AxioZoom V16 (Carl Zeiss, Brock Michelsen A/S, Denmark) (n= 29, per group). Trunk length, eye diameter, forebrain and full brain areas were measured using ImageJ software ^35^ and data were presented as mean ± SD. The 3 dpf zebrafish embryos were collected and fixed in 4% paraformaldehyde overnight at 4°C. Embryos were washed in 1X PBS, oriented and embedded in 5% agarose. The agarose block was dehydrated in alcohol solutions: 70% EtOH, overnight; 96% EtOH, overnight; 99% EtOH 3X 1h. The block was transferred into Xylene, 2X 1h and into paraffin overnight at 60**°**C. The block was embedded into fresh paraffin and 2 μm transverse sections were collected. Slides with the sections were dried overnight at 37**°**C followed by standard HE staining.

### *In situ* hybridization in zebrafish

The RT-PCR fragments of *neurod1* and *pcna* were amplified using the primers 5’-TAACCAGAGCATCCCATCAGC-3’/5’-GCGGCTTGACGTGAAAGATG-3’ and 5’-AGTGGACAGCACATCTGCAC-3’/5’-TGTGACCGTCTTGGACAGAG-3’ respectively and were cloned into the pCRII-TOPO vector (Invitrogen) for riboprobe synthesis. The plasmids were linearized and transcribed with SP6 or T7 RNA polymerases (Roche). Digoxygenin-labelled probes were used for single ISH and detected with NBT/BCIP (Roche). *In situ* hybridization was performed as described ^36^. Embryos were analysed on a Zeiss Axio Zoom V16 microscope.

### Statistical analysis

Data are presented as the mean ± SD. The sample size (n) indicates the experimental replicates from a single representative experiment; the results of the experiments were validated by independent repetitions. All experiments were carried out with n≥3. Statistical difference between groups (western blot band and fluorescence intensities) was determined by unpaired Student’s t test. n.s. indicates not significant, *p ≤ 0.05, **p ≤ 0.01, ***p ≤ 0.001, ****p ≤ 0.0001. All statistical tests were performed using GraphPad Prism version 8 or Windows Excel version 2010. Statistical analysis of haplotypes was performed using FASTLINK software. Logarithm of the odds (LOG) score was calculated at allele frequency (p=0.01), and disease frequency of 0.001.

## Supplementary Figures

**Supplementary Figure 1:**
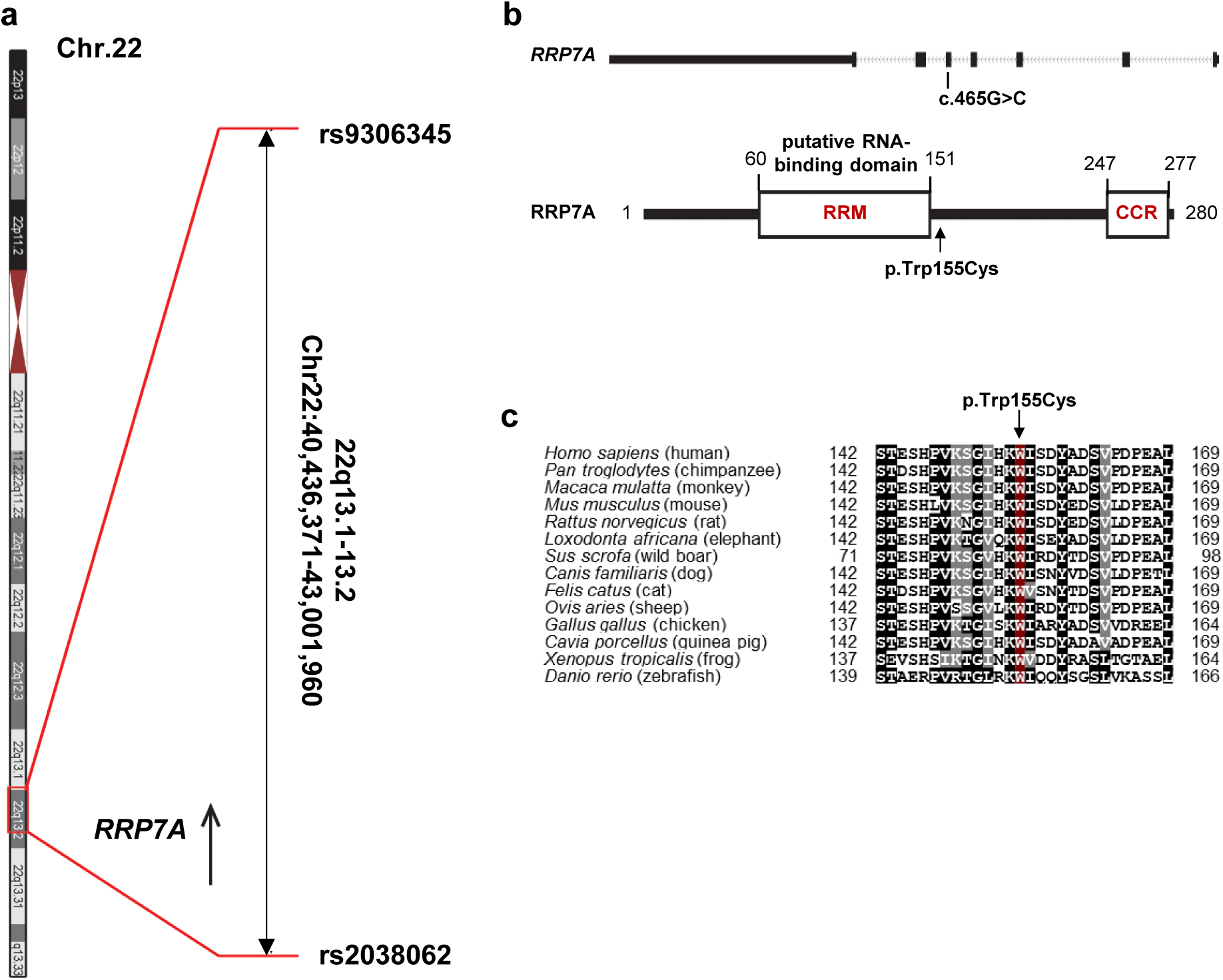
Identification of mutation in the *RRP7A* gene. **a**, Position of homozygous chromosomal region q13.1-13.2 at chromosome 22 marked by rs9306345 and rs2038062 (physical position 40,436,371-43,001,960, hg38). **b**, Multiple amino acid sequence alignment of RRP7A showing that W155 is highly conserved between different species. **c**, Schematic representation of the *RRP7A* gene (upper panel) showing site of mutation in exon 5. RRP7A protein domain prediction (lower panel) using SMART online tool predicted a RNA recognition motif (RRM; 60-151 amino acid), and a coiled coil region (CCR; 247-277 amino acid). The mutation p.W155C lies just outside the putative RNA binding domain (RRM). CCR: coiled-coiled region.

**Supplementary Figure 2:**
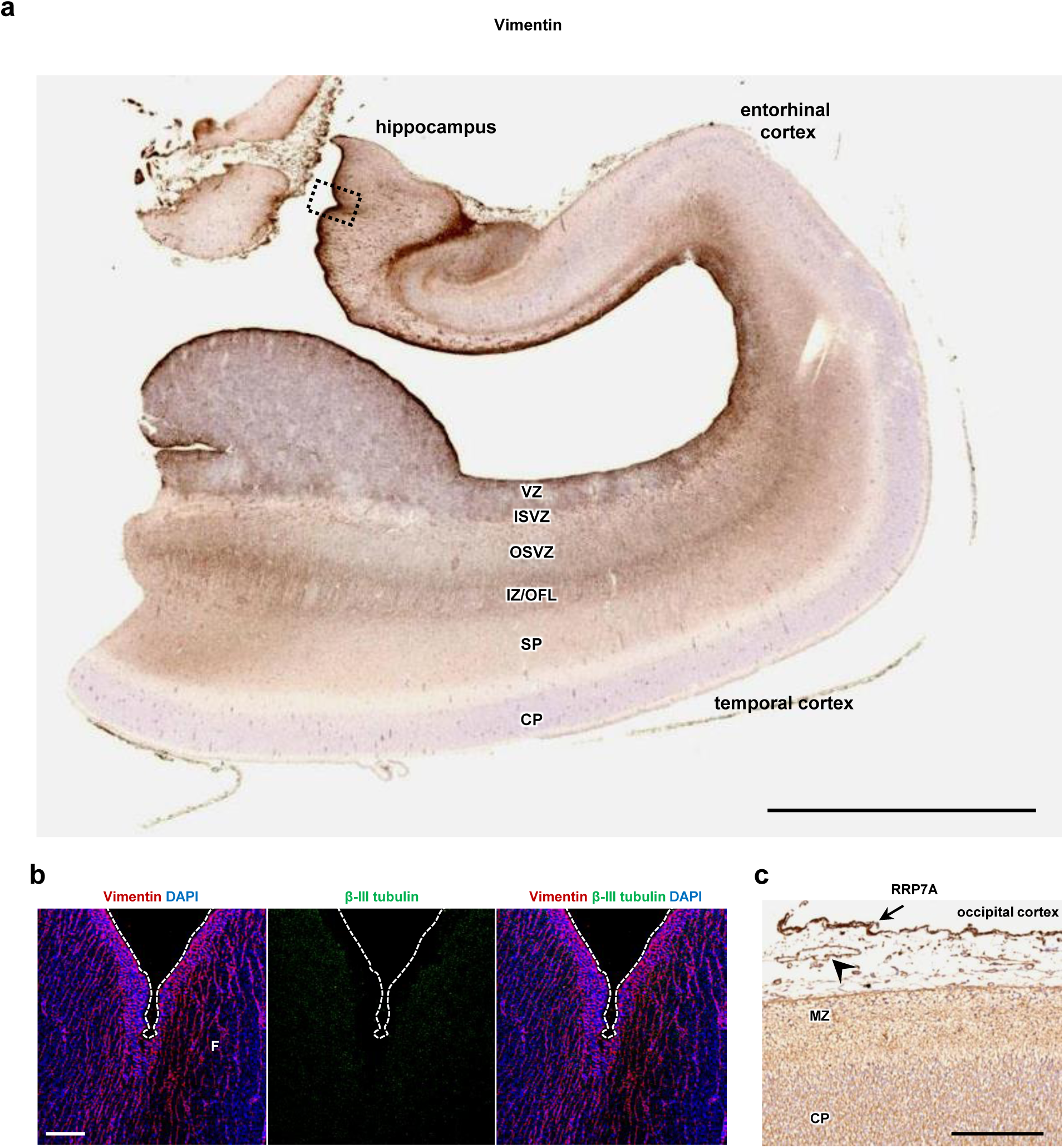
Expression of RRP7A in the human midgestation fetal brain aged 19 wpc. **a**, DAB staining of the entorhinal cortex and hippocampus for Vimentin. Scale bar, 5 mm. **b**, IFM analysis of the boxed area depicted in (a) showing lack of neurons (βIII-tubulin; *green*) at the outer surface along the hippocampus enriched in RGCs (Vimentin; *red*). Nuclei are stained with DAPI (*blue*). Scale bar, 0.1 mm. **c**, DAB staining of the occipital cortex depicting RRP7A localization to the meninges (arrow) and endothelial cells (arrow head). Abbreviations: VZ: ventricular zone, ISVZ: inner subventricular zone, OSVZ: outer subventricular zone, IZ/OFL: intermediate zone/outer fibre layer, SP: subplate, CP: cortical plate, F: Fimbria, MZ: marginal zone.

**Supplementary Figure 3:**
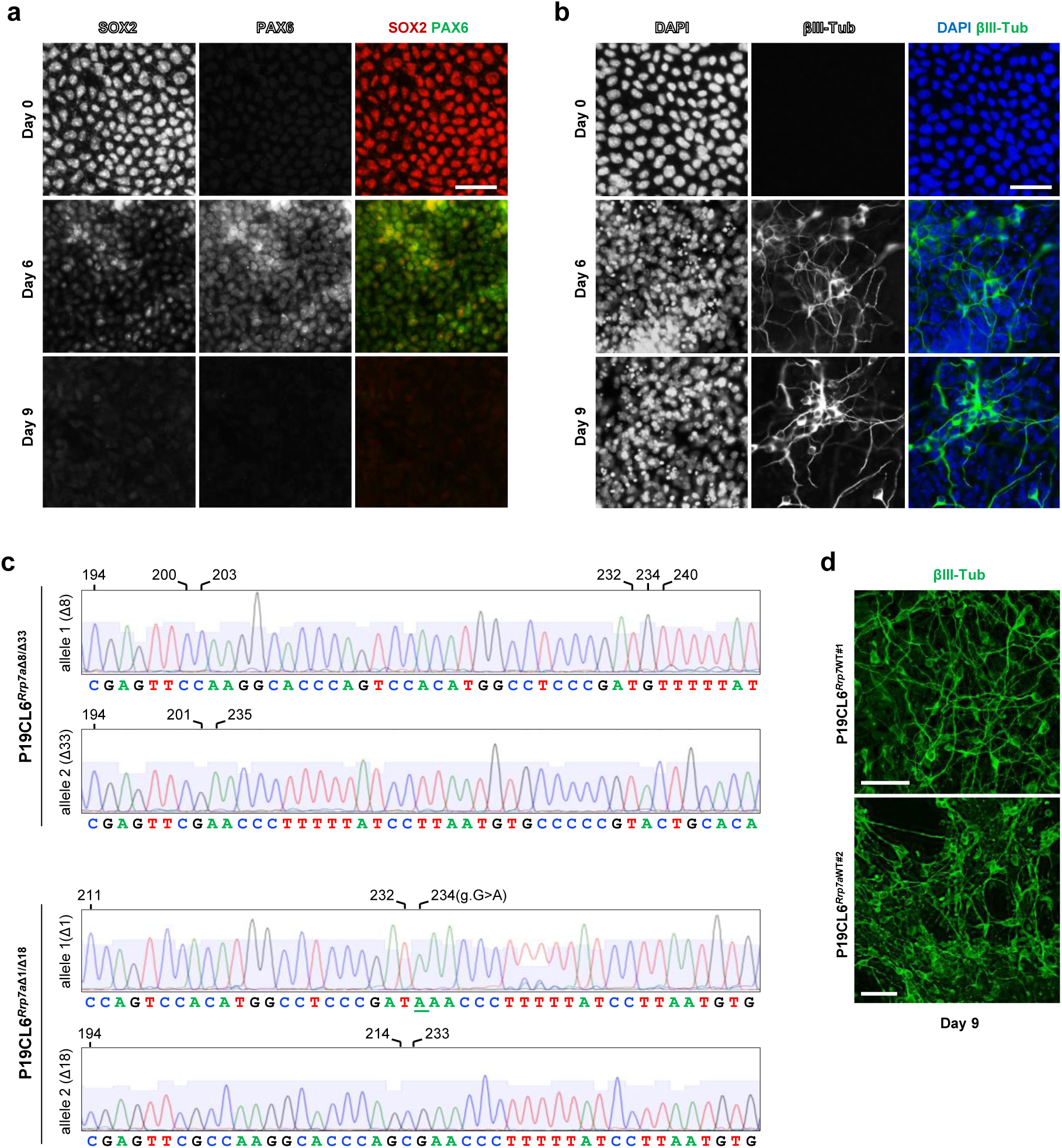
*In vitro* neurogenesis and Sanger sequencing of CRISPR clones in P19CL6 cells. **a**, IFM analysis of P19CL6 at days 0 (control), 3, 6, and 9 days of retinoic acid (RA) treatment, showing expression of the stem cell marker SOX2 (*red*) and the neurogenesis-controlling factor PAX6 (*green*). Scale bar, 40 mm. **b**, IFM analysis of P19CL6 cells at equivalent days of RA treatment, showing expression of ßIII-tubulin (*green*). Nuclei are stained with DAPI (*blue*). Scale bar, 40 mm. **c**, Sanger sequencing of mutant clones P19CL6^*Rrp7a*Δ8/Δ33^ and P19CL6^*Rrp7a*Δ1/Δ18^. Deletions are indicated by base numbers above diagrams. **d**, IFM analysis showing that WT CRISPR clones form neurons within 9 days of RA treatment ßIII-tubulin (*green*), DAPI (*blue*). Scale bar, 40 µm.

**Supplementary Figure 4:**
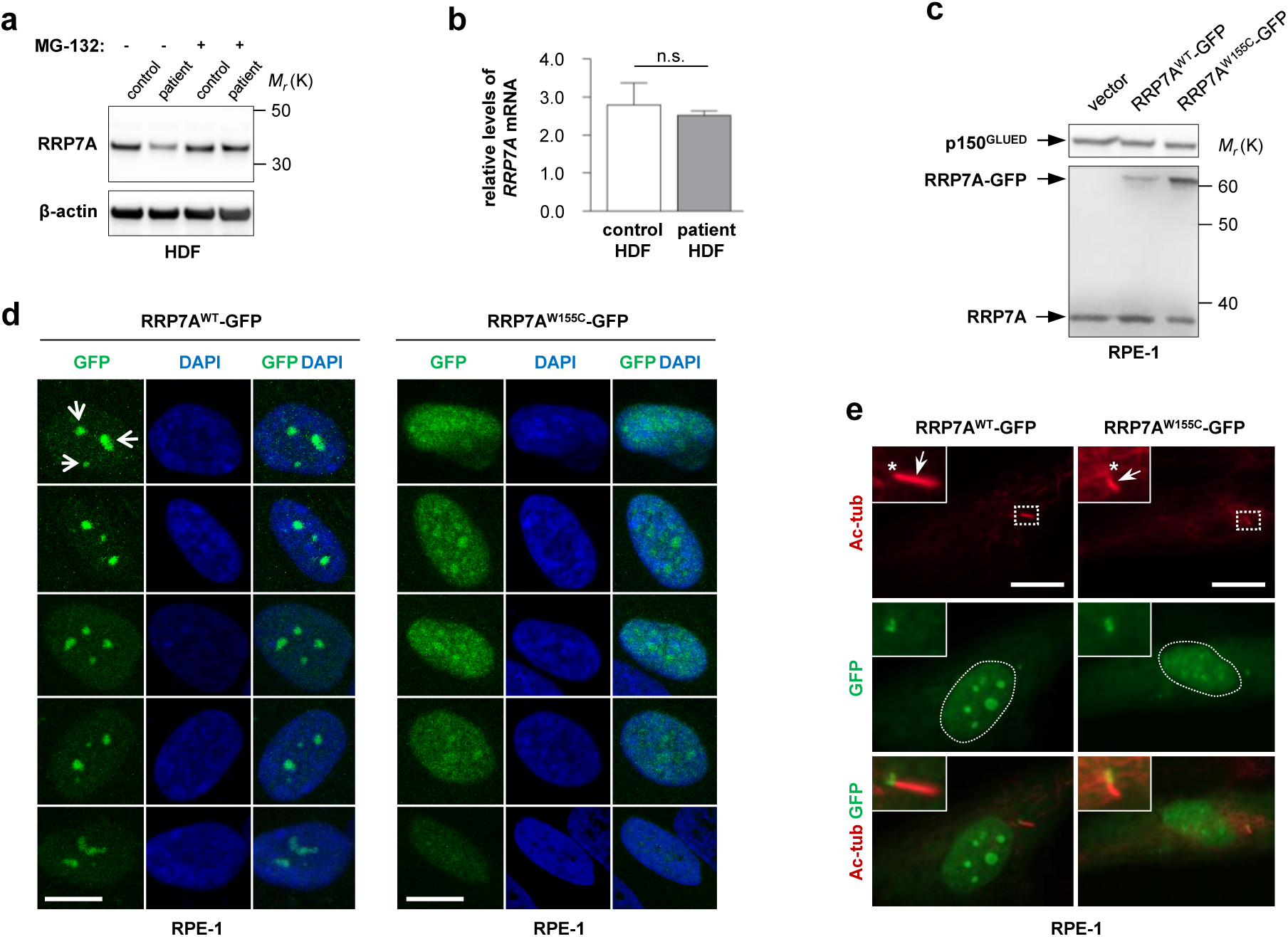
Expression and localization of RRP7A and RRP7A constructs in HDFs and hTERT RPE-1 cells. **a**, WB analysis of RRP7A expression in control and patient HDFs in the absence and in the presence of the proteasome inhibitor MG-132. β-actin: loading control. **b**, Quantification of relative levels of *RRP7A* mRNA detected by qRT-PCR in control and patient HDFs (n=3). **c**, WB analysis of hTERT RPE-1 (RPE-1) cells transiently transfected with empty vector or vectors expressing GFP-tagged constructs of either WT (RRP7A^WT-GFP^) or patient mutant (RRP7A^W155C-GFP^) versions of RRP7A. p150^GLUED^: loading control. **d**, IFM analysis of localization of RRP7A^WT-GFP^ or RRP7A^W155C-GFP^ (GFP, *green*) to nucleoli (arrow heads) in serum supplemented cultures of RPE-1 cells. Nuclei are stained with DAPI (*blue*). Scale bar 10 µm. **e**, IFM analysis of localization of RRP7A^WT-GFP^ or RRP7A^W155C-^ GFP (GFP, *green*) to nucleoli and primary cilia (closed arrows, Acetylated α-tubulin, Ac-tub, *red*) in serum-depleted cultures of RPE-1 cells. Asterisks indicate the ciliary base and the dotted lines outline the nucleus. Scale bar, 10 µm.

**Supplementary Figure 5:**
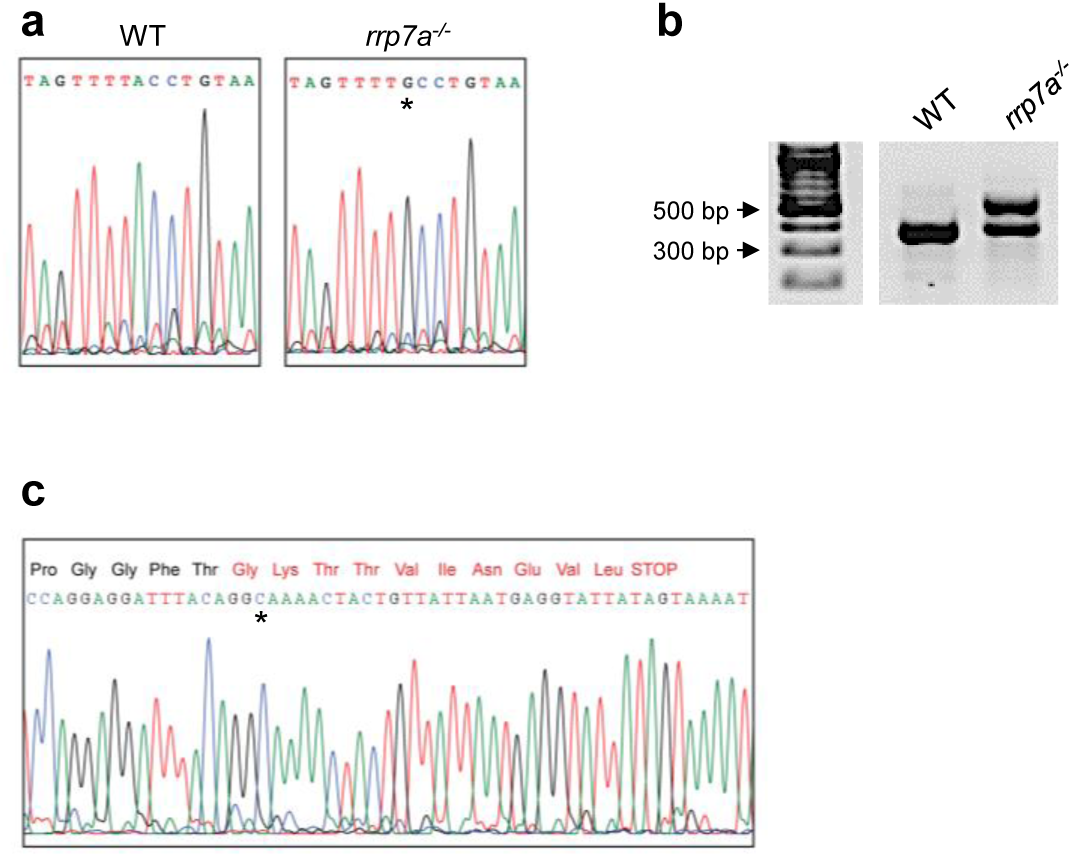
Zebrafish *rrp7a* mutants. **a**, Reverse-complement *rrp7a* DNA sequence of WT and homozygous mutant embryos. Mutants carry a point mutation in the donor-splice site of exon 1 in *rrp7a* (ivs1 +2 T>C, NM_001017579), leading to aberrant splicing of *rrp7a* mRNA, with inclusion of 191 bp of intronic sequence. The inclusion of intronic sequence leads to a shift in the reading frame, inclusion of 10 abnormal amino acid residues at position 18-27 followed by a premature stop codon. The deduced mutant protein is truncated C-terminal to the RRM and contains 27 amino acid residues in total. The ivs1 + 2 T>C mutation is marked with *. **b**, RT-PCR fragments from *rrp7a* mRNA extracted from WT and homozygous mutant embryos. **c**, DNA sequence of aberrantly spliced *rrp7a* mRNA (upper right band in B). Ivs1 + 2 T>C mutation is marked with an asterisk. Abnormal amino residues are shown in red.

## Supplementary Tables

**Supplementary Table 1:** Clinical phenotypes of affected individuals. Clinical characteristics of eight effected family members. HC: Head Circumference SD: Standard deviations from age-adjusted mean.

| Patient | Age | Sex | HC (cm), SD | Intellectual disability | Speech impairment |
| --- | --- | --- | --- | --- | --- |
| IV.3 | 30 | Male | 44, -8 SD | Moderate | No |
| V.4 | 15 | Female | 45, -7 SD | Mild | NO |
| V.6 | 12 | Female | 42.5, -8 SD | Mild | YES |
| V.8 | 8 | Female | 41, -7 SD | Moderate | NO |
| V.10 | 10 | Female | 42.5, -6 SD | Mild | NO |
| V.12 | 27 | Male | 47, -6 SD | Moderate | YES |
| V.14 | 14 | Female | 45.5, -7 SD | Severe | YES |
| V.15 | 11 | Female | 43.5, -6 SD | Mild | YES |

**Supplementary Table 2:** List of primary antibodies. Abbreviations: WB: western blotting, IFM: immunofluorescence microscopy, DAB: 3,3’-diaminobenzidine.

| Primary antibody | Species | Supplier | Dilution |
| --- | --- | --- | --- |
| Acetylated α-tubulin | Mouse | Sigma-Aldrich, T7451 | 1:2000 (IFM, DAB) |
| ARL13B | Rabbit | Proteintech, 17711-1-AP | 1:600 (IFM) |
| βIII-tubulin | Rabbit | AbCam, AB18207 | 1:200 (WB)<br>1:2000 (IFM, DAB) |
| GFAP | Rabbit | DAKO, Z0334 | 1:1000 (DAB) |
| MAP2 | Rabbit | Abcam, AB32454 | 1:2000 (WB) |
| MAP2 | Chicken | Abcam, AB5392 | 1:1000 (IFM) |
| p150-GLUED | Mouse | BD Bioscience, 610474 | 1:1000 (WB)<br>1:500 (IFM) |
| PAX6 | Rabbit | Abcam, AB5790 | 1:300 (WB)<br>1:300 (IFM) |
| Pericentrin-2 (C16) | Goat | Santa Cruz, sc-28145 | 1:300 (IFM) |
| p-RB | Rabbit | Cell Signaling, 9308s | 1:300 (WB)<br>1:500 (IFM) |
| RRP7A | Rabbit | AbCam, AB185225 | 1:2000 (WB) |
| RRP7A | Rabbit | Sigma, HPA001586 | 1:500 (IFM, DAB) |
| RRP7A | Mouse | Santa Cruz, SC377210 | 1:500 (IFM) |
| SOX2 | Mouse | R&D system, MAB2018 | 1:400 (IFM) |
| Vimentin | Chicken | Thermo Fischer, PA1-16759 | 1:200 (IFM) |
| Vimentin | Mouse | Dako, M0725 | 1:100 (DAB) |

**Supplementary Table 3:**
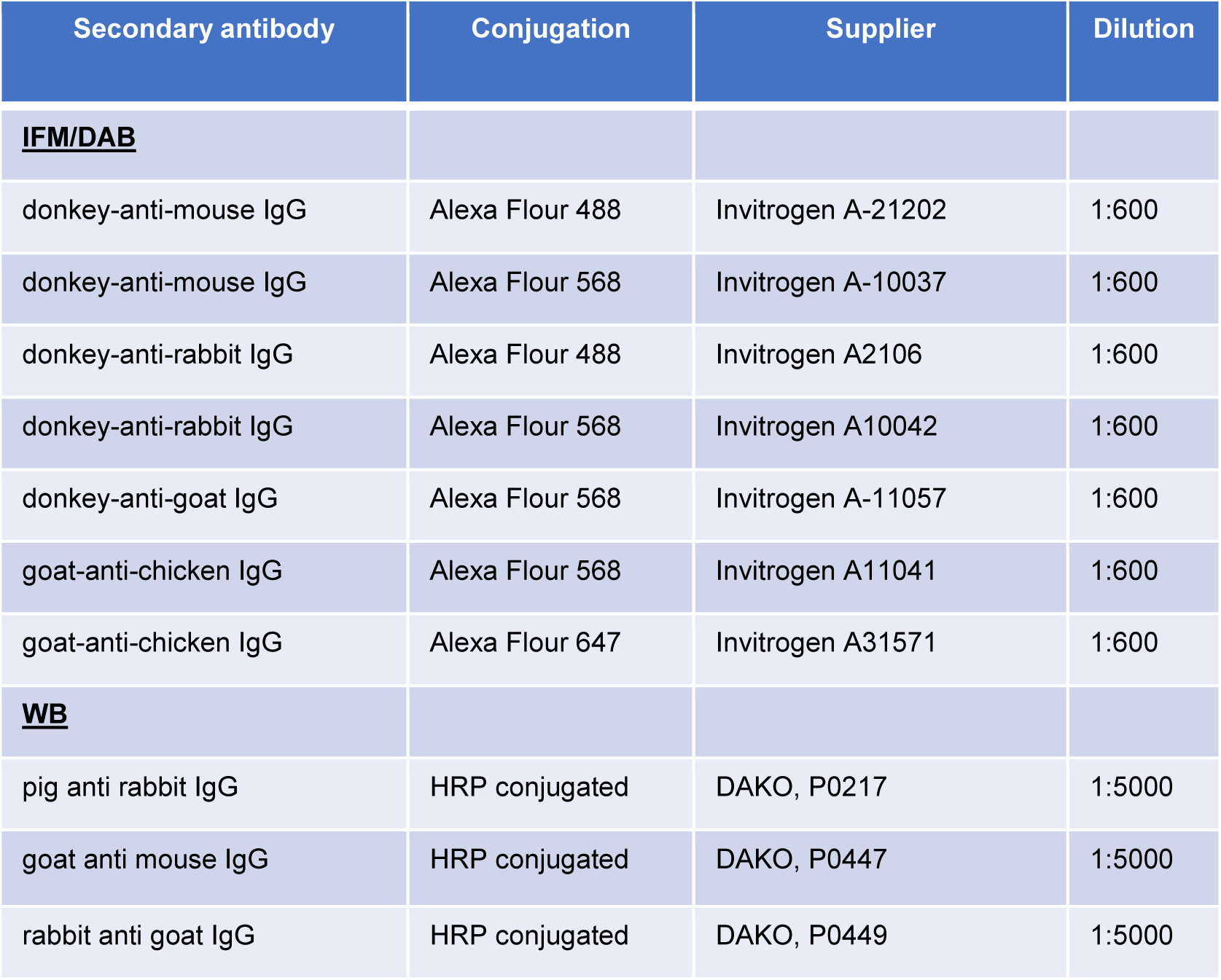
List of secondary antibodies. Abbreviations: WB: western blotting, IFM: immunofluorescence microscopy, DAB: 3,3’-diaminobenzidine.

